# Density dependent interactions during growth mask the effect of interactions during flowering in pollinator-sharing annual plants

**DOI:** 10.1101/2021.03.31.437969

**Authors:** Aubrie R. M. James, Monica A Geber

## Abstract

Density dependent interactions are fundamental to community ecology, but studies often reduce the complex nature of species interactions. In plant ecology, interactions during vegetative growth and flowering are often considered separately, though both can affect reproductive output. Here we use communities of annual flowering plants in the genus *Clarkia* to ask how interactions during growth and flowering contribute to density dependence in plant seed production, and if pollinator behaviors explain apparent patterns in plant interactions during flowering. We measure seed set (seed number per ovule) and total fecundity (whole-plant seed production) of *Clarkia* focal plants in experimental interaction plots with the effect of pollinators experimentally removed through supplemental pollination or retained. We also observe pollinator behaviors in the plots and experimental arrays to document pollinator preference, constancy and joint attraction. During flowering, pollinators significantly changed the density dependent effects of *Clarkia* interactions on seed set in 31% of species interactions, and these changes corresponded to pollinator behaviors. Total fecundity, however, did not depend on interactions between *Clarkia*; instead, earlier-flowering, non-*Clarkia* forbs limited total fecundity. Our study shows that interactions during vegetative growth can preclude the effect of pollinator-mediated interactions on fecundity by limiting potential reproductive output. Simultaneously studying different types of species interactions allows for understanding the contingency of ecological outcomes.

## Introduction

Density dependent interactions shape population dynamics and community structure. In communities, co-occurring species are typically engaged in multiple types of interactions within and between trophic levels that can affect density dependence (e.g., plant-plant resource competition and pollinator-mediated plant interactions). Typically, studies aimed at quantifying the strength and magnitude of density dependent interactions either estimate the combined effects of multiple interaction types on performance or focus on a single interaction type (Losapio et al. 2019). For example, studies in plant community ecology estimate the effects of species interactions on coexistence (e.g. Angert et al. 2009, Godoy et al. 2014, Kraft et al. 2015, Bimler et al. 2018, Wainwright et al. 2019), but do not parse the contribution of plant-plant interactions during vegetative growth from the effects of pollinator-mediated interactions during flowering (but see Lanuza et al. 2018 for an example of how variation in pollination affects pairwise coexistence dynamics). Conversely, recent work in pollination ecology demonstrates that pollinator-mediated plant interactions are key to understanding seed production and density dependence (Lázaro et al. 2014, Benadi & Pauw 2018, Bergamo et al. 2020, Opedal & Hegland 2020, Phillips et al. 2020), but rarely do these studies investigate how interactions during vegetative growth also contribute to seed production. On one hand, the ‘combined effects’ approach cannot identify how interaction types combine to determine overall patterns in density dependence, which lowers its predictive value. The ‘single interaction’ approach, on the other hand, assumes that only one interaction type is relevant for reproductive output and therefore may overestimate its importance, which lowers the explanatory value. One way to unite these two approaches in plant ecology is to quantify density dependence arising from interactions during growth and flowering and determine their relative contributions to seed production. This should illuminate the ways in which interactions during these two life stages might combine to affect plant species’ performance, population persistence, and coexistence (Bartomeus et al. 2021).

For plant species that grow together and share pollinators, density dependent interactions during vegetative growth determine plant survivorship, size, and potential fecundity (e.g., flower and ovule production). Interactions during the reproductive stage determine whether flowers are pollinated and produce seed. Whereas the mechanisms driving interactions during vegetative growth have been widely explored (e.g. Dybzinski & Tilman 2007), pollinator-mediated interactions during flowering are less understood. Two aspects of pollinator foraging behavior, pollinator preference and constancy, have the potential to explain the effect of pollinator mediated interactions on seed set. Preference reflects a pollinator’s choice in visiting one plant species over others when all are available and can introduce or intensify competition between plant species by affecting the quantity of conspecific pollen that plants receive. Conspecific pollen receipt and seed set for a particular plant should decline with declining pollinator preference (Pauw 2012, Song & Feldman 2014, Benadi 2015). Constancy is the tendency of pollinators to visit one species of plant in a foraging bout. If pollinators are inconstant, they can introduce incompatible, heterospecific pollen to co-flowering plants. Heterospecific pollen transfer is widespread in plant communities and is a form of reproductive interference between plant species that decreases seed set (Morales & Traveset 2008, Arceo-Gómez et al. 2016, Arceo-Gómez et al. 2019, Moreira-Hernández & Muchhala 2019). Plant interactions via shared pollinators can also be facilitative. Patches of many co-flowering plants may attract more pollinators than when growing alone, which can increase pollinator visitation rates and thus pollination service; this phenomenon is known as joint attraction (Kunin & Iwasa 1996, Moeller 2004, Feldman et al. 2004, Bizecki Robson 2013). The potential for facilitation via joint attraction is contingent on low interspecific pollen transfer, and thus high pollinator constancy. Lastly, because different pollinator taxa exhibit different degrees of preference and constancy (Waser et al. 1996; e.g. Eckhart et al. 2006, Barrios et al. 2016), the composition of the pollinator community will also modulate the extent to which pollinator-mediated interactions affect plant reproduction.

Studies of in natural plant-pollinator communities (Lanuza et al. 2018) and small mesocosms of plant and pollinator species (Bartomeus et al. 2021) confirm the importance of examining plant-plant interactions during both vegetative growth and flowering. However, no experimental study has quantified the relationship between density dependence arising from interactions during vegetative growth versus flowering, nor demonstrated the link between pollinator-mediated interactions during flowering and pollinator foraging behavior. In this study, we do both in a group of four sympatric and co-flowering winter annuals in the genus *Clarkia* (Onagraceae).

The four *Clarkia* species commonly co-occur throughout their range of geographic overlap (Eisen & Geber 2018) and share bee pollinators specialized on the genus (MacSwain et al. 1973, Moeller 2005, Singh 2014), suggesting that they interact during both vegetative growth and flowering. Furthermore, because *Clarkia* species flower late in the season after almost all other annual plants in the area have senesced, the main interaction during flowering is through pollinators that they share with each other (MacSwain et al. 1973).

To investigate the separate and combined effects of density dependent interactions during growth and flowering on plant reproductive success, we use experimental pairwise interaction plots of sown *Clarkia* and pollen supplementation via hand pollination (Figure 1). In the pairwise interaction plots, interactions during growth affect plant size, and hence reproductive potential, while interactions via shared pollinators during flowering affect pollination and seed set. The pollen supplementation alleviates any effect of pollen limitation to seed set, allowing the effect of interactions during growth to be separated from the effect of pollinator-mediated interactions during flowering. We also investigate the link between pollinator behavior and *Clarkia* interactions during flowering using pollinator observations in our experimental interaction plots, and in experimental floral arrays, to measure pollinator preference, constancy, and visitation rate in response to floral density.

**Figure 1.**
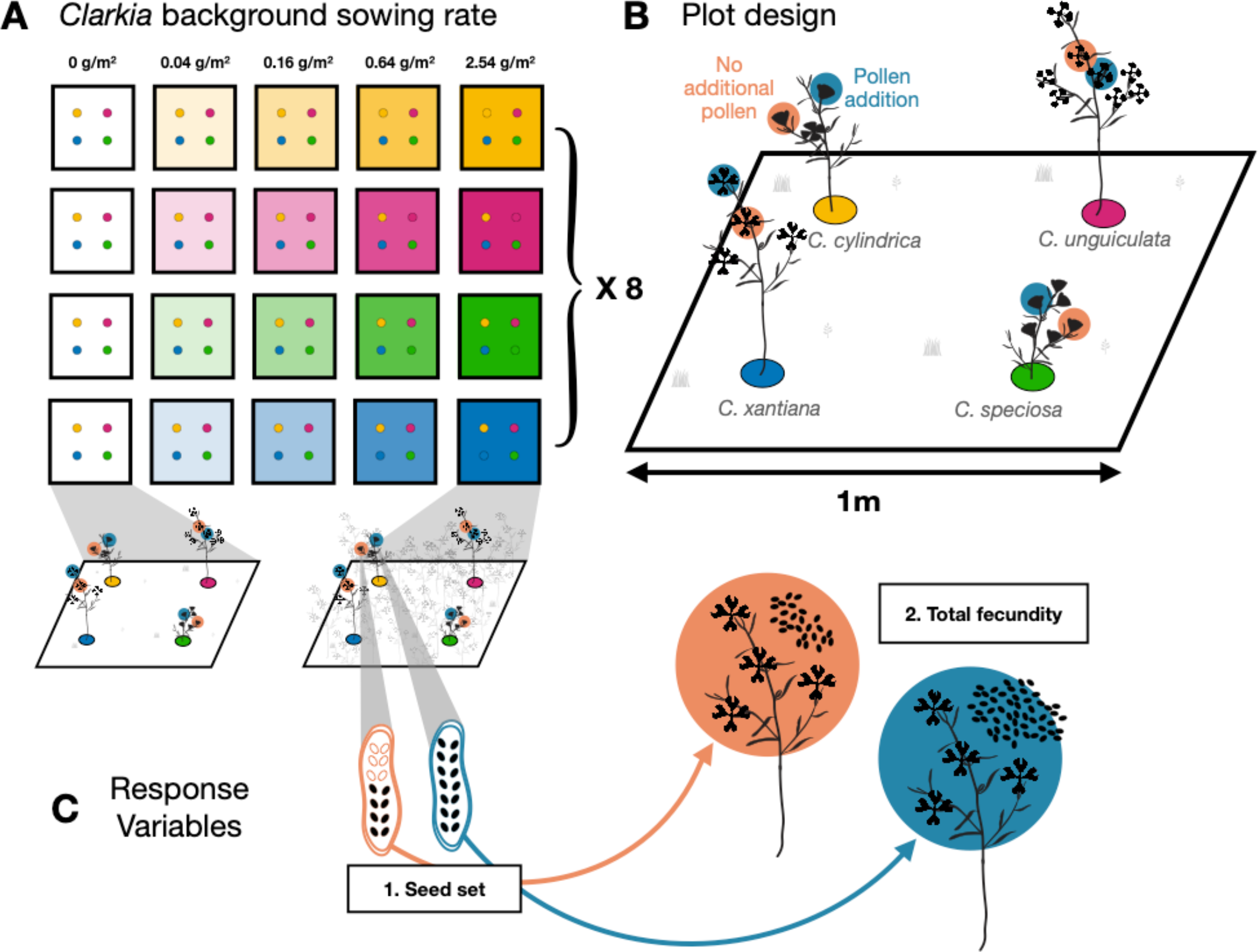
Schematic of the pairwise interaction plot experiment. A. In experimental 1m x 1m plots, seeds of one *Clarkia* species (yellow: *C. cylindrica*, green: *C. speciosa*, pink: *C. unguiculata*, blue: *C. xantiana*) were sown in the background at a range of seeding rates from 0g/m2 to 2.54g/m2; 20 seeds of a focal *Clarkia* species were sown in one of the four cabone rings secured in the four corners of the plot. Following germination, focal seedlings in the cabone rings were thinned to a single individual. The schematic shows the range of background treatments but does not depict the actual arrangement of plots in the field. Four plots with a given background species and varying background seeding densities were grouped into blocks. There were 8 replicate plots of each background species x background seeding density. B. Depiction of pollination treatments on focal plants. In each plot, two pollination treatments were applied to all focal plants that had 2 or more flowers. One flower was assigned as the control and received no additional pollen while the second flower received additional pollen from hand pollination. Pollen was collected from the appropriate *Clarkia* species from plants growing at the site but outside of experimental plots. C. Two response variables were measured on each focal plants 1) seed set equal to the ratio of filled seed to total ovules (unfertilized ovules, aborted ovules and filled seed) in fruits from control and pollen-addition flowers; 2) total fecundity equal to the product of fruit number, average total ovule number per fruit, and average seeds per fruit in the two different pollination treatments.

## Methods

### Natural history

*Clarkia* is a genus of winter annual plants which germinate in late winter and grow over the course of the spring. All *Clarkia* species are self-compatible, but 2/3 of the species are predominantly outcrossing due to pronounced spatial separation (herkogamy) of anthers and stigmas within flowers and temporal separation (protandry) between anther dehiscence and stigma maturation (Lewis & Lewis 1955). All four *Clarkia* taxa in this study (*C. cylindrica* ssp. *clavicarpa* (Jeps.) Lewis & Lewis*, C. speciosa* ssp. *polyantha* Lewis & Lewis*, C. unguiculata* Lindl.*, C. xantiana* ssp. *xantiana* A. Gray, family Onagraceae) exhibit pronounced herkogamy and protandry, and in the two taxa where outcrossing rates have been measured, outcrossing rates range from 0.53-0.95 in *C. xantiana* ssp. *xantiana* (Moeller et al. 2012; Hove et al. 2016) and 0.80-0.94 in *C. unguiculata* (Hove et al. 2016). In the remainder of the paper, we drop subspecies designations and refer to the four taxa as species.

The four *Clarkia* species are sympatric in the southernmost portion of the Sierra Nevada Mountain range. This region and our study are on Tübatulabal land known commonly as the Kern River Canyon (KRC; Kern County, California, USA). We pay our respects to the Tübatulabal people and recognize their past and ongoing connection to the land and its ecology. The *Clarkia* in the KRC co-occur in communities of varying congeneric composition (from one to four *Clarkia* species) embedded in low-elevation grassland, chaparral, and oak woodlands. In their range of sympatry, *Clarkia* begin to flower in early May, and flowering lasts until mid- to late June. All four species overlap in flowering time for about two weeks in late May and share a group of bee pollinators specialized on the genus rather than any one species (MacSwain et al. 1973, Singh 2014). Pollen limitation to reproduction has been found in *C. xantiana* (Moeller et al. 2012, Hove et al. 2016) and *C. cylindrica* (Eisen et al. 2020), but not *C. unguiculata* (Hove et al. 2016). Where present, pollen limitation is variable across sites and years (Moeller et al. 2012, Hove et al. 2016, Eisen et al. 2020). Research has shown that at least one species, *C. xantiana*, has higher seed set when it grows in multi-species *Clarkia* communities than at sites where it occurs alone, and that this effect is explained by increased visitation from shared pollinators at multi-species sites (Moeller 2004).

We use two different experiments to understand how pollinator-mediated interactions factor into interactions between *Clarkia*. First, we use an *in-situ* common garden experiment to determine (1) how focal *Clarkia* seed production and total fecundity respond to variation in neighborhood density; (2) how pollinator-mediated interactions contributed to patterns in observed density-dependence; and (3) pollinator preference and joint attraction. Because pollinator constancy was impossible to estimate in pairwise interaction plots due to low species richness, we additionally recorded pollinator visitation in multi-species experimental arrays of *Clarkia* plants to estimate pollinator constancy.

### Experimental Design

#### Pairwise Interaction Plots

Interactions during growth and flowering

To determine the combined effects of *Clarkia* growing and flowering together on reproductive success, we used an experimental design closely modeled after work done to estimate competitive interactions in annual plant communities (Godoy & Levine 2014, Godoy et al. 2014, Kraft et al. 2015, Wainwright et al. 2019). In short, seeds of one *Clarkia* species were sown into a plot as a competitive background, and then seeds of the same and different *Clarkia* species were sown as focal plants (Figure 1A, B). The identity of the background *Clarkia* species, and the density of background *Clarkia* varied across plots. This allowed us to estimate the effect of increasing density of background *Clarkia* on focal plant fecundity for each pair of focal species and background species.

In the pairwise interaction plots, we measured two metrics of reproductive success on focal *Clarkia* plants in response to neighborhood density. First, we measured seed set as the success of converting ovules in a fruit into seeds (number of seeds per ovule or the proportion of ovules that result in filled seed) (Figure 1C). This metric is determined by interactions that occur during pollination. Second, we measured total fecundity, or the total number of filled seeds produced by a plant (Figure 1C), determined by interactions over the entire lifespan of the plant. Total fecundity is the product of the total number of ovules a plant produces, which is determined during growth and represents a plant’s maximum potential fecundity, and seed set, as described above. In our experiment, both seed set and total fecundity are potentially determined by the density and identity of background sown *Clarkia*.

To remove the effects of pollinator-mediated pollen limitation to seed set, we added conspecific pollen to one flower on each focal plant (Figure 1B). Comparison of seed set in the pollen supplemented fruit, whose flower received additional pollen, to seed set in a control fruit, whose flower did not receive additional pollen quantifies the effect of pollinator-mediated pollen limitation to seed set (Figure 1C). We refer to these two fruits as “pollen-supplemented” and “control” fruits in the rest of the paper. Total fecundity calculated from the seed set of control fruits reflects the combined effects of plant-plant interactions during growth and, via pollinators, during flowering on plant reproductive success, while total fecundity calculated from the seed set of pollen-supplemented fruits reflects only the effects of plant-plant interactions during growth on reproductive success (Figure 1C).

In late June through early July 2016, we set up 160, 1X1 meter experimental plots in a community where all four species naturally occur in the Kern River Canyon (35°31’45N, 118°39’36W; Area = 0.57ha). We arranged plots in blocks of four, where plots were separated by ∼1/4 meters, and blocks were separated by ∼1/2 meter. We cleared plots of all naturally occurring *Clarkia* to prevent natural seed rain into the plots but did not disturb the soil. We also left non-*Clarkia* forbs and grasses in place for two reasons: because it was infeasible to weed them, and because grasses and forbs make up a substantial component of the vegetation; removing them could disrupt interactions among *Clarkia*, and our intention was to study interactions among *Clarkia* species in as natural a setting as possible (Table of common grass and forb species in Table S1).

*Clarkia* seeds for the experimental sowing were collected at the study site as well as nearby sites in the Kern River Canyon at the end of the 2016 growing season (late June). We used a hand deawner (Hoffman Manufacturing, Inc, Fogelsville PA, USA) to break open the fruits and extract seed, and a seed blower to separate seed from fruit walls. We sowed the seed into experimental plots in October 2016 before the onset of winter rains that trigger germination.

Each block was haphazardly assigned a background *Clarkia* species, and each plot within the block was haphazardly assigned one of five seeding densities (0, 0.04, 0.16, 0.64, and 2.56 g/m^2^). For a given background seeding density, we chose to plant the same mass of seed because the four species differ in seed size, and seed size correlates with adult plant size. Had we planted equal seed numbers for a given seeding density, the background biomass, and hence competitive effect, of large seeded species (*C. unguiculata* and *C. xantiana*) would have been greater than that of the smaller seeded species (*C. cylindrica* and *C. speciosa*).

Background seeding densities were chosen based on previous estimates of naturally occurring seed production at the field site (James, unpublished data). For *C. xantiana*, with an average seed mass of 0.025 mg (Gould et al. 2014), the five seeding densities corresponded to 0, 1600, 6400, 25600, and 102400 seeds per m^2^. In total, our design yielded 20 background species X density treatments, with eight plot replicates per treatment.

To establish focal plants of each species on which to measure reproductive output, we planted twenty focal seeds into one of four designated locations (one for each species) in each plot. The locations were delimited using 2.5cm radius cabone craft rings (CSS Industries, Inc., PA, USA) secured in the ground with two 3.8cm fence staples. In the spring of 2017, we thinned the focal seedlings in the cabone rings to one focal *Clarkia* plant. To prevent bias in selecting a focal seedling, we left the seedling that was closest to the center of the ring and removed the rest. In this same period, we counted the number of seedlings of background *Clarkia* species that had germinated and the number of non-target grasses and forbs growing within a 5cm radius from the focal seedling. We also weeded any seedlings of non-target *Clarkia* species in the background of all plots.

During the two-week flowering stage in late May 2017 we haphazardly assigned one flower as the control to receive no additional pollen, and another flower to receive additional conspecific pollen on all focal plants with ≥ 2 flowers. We selected pairs of flowers that were at a similar reproductive stage so that they were exposed to similar environments during pollination. To add conspecific pollen to the pollen addition flowers, we harvested mature, dehiscing anthers from 3-4 flowers from naturally occurring conspecific *Clarkia* growing nearby by 8 am before pollinators began foraging for the day. The pollen was applied to the freshly opened and receptive stigmas on the selected flower of a focal plant immediately after harvesting the anthers. Applying pollen to freshly opened stigmas ensured that they had not received any con- or heterospecific pollen prior to applying our treatment. Application of pollen in this manner coats the stigma with >300 pollen grains, well in excess of the number of ovules (20-100) in each ovary (M.A. Geber, pers. obs.).

We counted the number of fruits on each focal plant and collected the treatment fruits at the end of the growing season. We also counted the number of filled seeds, aborted seeds, and unfertilized ovules in each treatment fruit. We harvested all background *Clarkia* in each plot and weighed the above ground biomass after drying plants at 65 C for 48 hours.

### Pollinator visitation in the interaction plots

During the co-flowering period in 2017, we observed bee visitation in the pairwise interaction plots to quantify pollinator preferences and joint attraction (Figure S1). Due to time limitation and the fact that some plots did not have surviving focal plants, not all 1m^2^ plots were observed; however, at least one plot from each block was observed during the co-flowering period.

Observations were made in ten-minute periods. Bees entering plots were identified to genus or species and tracked as they made visits to *Clarkia* flowers. After the 10-minute observation period, we counted and recorded the number and identity of open *Clarkia* flowers in the plot.

### Pollinator Arrays to assess constancy

We assembled cut-stem arrays of the four *Clarkia* species to observe pollinator behavior during the 2015 and 2016 *Clarkia* growing seasons. We cleared a 5 x 5m patch of all naturally occurring *Clarkia* in the middle of the experimental site in late April before *Clarkia* plants had started flowering. We constructed a grid with nodes at every 0.5m increment for a total of 100 nodes and buried a floral water pick at each node to fill with water and hold *Clarkia* stems.

Each array contained equal numbers of cut stems of all four *Clarkia* species planted at one of four densities (in 2015: 10, 20, 40, or 100 stems; in 2016: 12, 24, 64, or 100 *Clarkia* stems) (Table S1). This design allowed us to observe pollinator constancy across a range of plant densities, holding diversity relatively constant. Because different bee species forage at different times of day (Hart & Eckhart 2010), we ran most treatments in the morning and afternoon on different days. For each array run, we gathered *Clarkia* stems before 8am to assemble in the grid. We arranged *Clarkia* stems in the grid according to a randomly generated map of a particular treatment, where locations in the grid were either left ‘blank’ or received one *Clarkia* stem of a prescribed *Clarkia* species depending on the treatment density.

During the array runs, we identified and monitored bees as they entered and foraged in arrays. When a bee entered the array and landed on a flower, one person called out the grid location of the flower/bee, identified the bee to species or genus, and tracked the bee’s movements until it left the array, while another person recorded. At the end of each array run, we counted the number of open flowers of each species and then disposed of wilted *Clarkia* stems. Rarely, we re-used stems if they still had visibly turgid petals and leaves, though we never retained stems for more than two runs. Array treatments lasted 40 or 60 minutes; many treatments in 2016 were stopped at 40 minutes due to wilting. We did not run the experiment on days when the predicted peak temperature was below 21°C or wind was stronger than a light breeze.

### Statistical Analysis

All calculations and statistical analyses were performed in R (R version 3.5.2, R Core Team, 2018). We used two different analyses to understand the effects of density dependent interactions on *Clarkia* seed production. We first investigated density dependent interactions in the flowering stage by determining how *Clarkia* densities affected seed set (the number of seeds per ovule in a fruit), and then investigated the density dependent effects of the neighborhood on *Clarkia* total fecundity using glmms. We then analyzed the three pollinator foraging behaviors that contribute to density dependence during the flowering stage.

### Seed set

For each fruit, we defined ‘successes’ as filled seeds, and ‘failures’ as the sum of unfertilized ovules and aborted seeds. We built a set of candidate models to determine which combination of four fixed effects factors best explained variation in seed set – the pollination treatment (pollen supplemented vs. control), the focal *Clarkia* species identity, the background *Clarkia* species identity, and the biomass of background *Clarkia* in the plot. We did not include non- *Clarkia* grasses or forbs in this analysis because they flower before *Clarkia* during the experiment, and therefore would not affect the pollination interactions of *Clarkia*. We also used one random effect, the identity of the focal plant, in all models. We did not include random effects for plot nested within block or for block because models with these effects were unidentifiable. To avoid model dredging, we built a set of models relevant to our questions and hypotheses with which to perform model selection (Table S3). We fit the data to all candidate models using the lme4 package (Bates et al. 2015) and selected the model with the lowest AICc (the best-fit model) for ecological inference. The most complex model included an interaction of all four fixed effects and represents our prediction that successful seed set depends on the density dependent effect of the background *Clarkia* on the focal *Clarkia* and varies according to pollination treatment.

After identifying the best-fit model, we used estimated marginal means to determine the predicted density dependent effect of background *Clarkia* on focal *Clarkia* seed set, with and without pollen addition (package emmeans, Lenth 2019). Here, the effect of pollinators on the interaction is the change in the density dependent effect of background *Clarkia* on focal *Clarkia* seed set between the two pollination treatments (pollen addition and no pollen addition). If pollinators change the strength or direction of the interaction, then the effect of increasing background *Clarkia* biomass will be significantly different between the two pollination treatments. We also tested if the estimated density dependent effect of background *Clarkia* on focal *Clarkia* was significantly different from zero for both pollination treatments to examine if and when pairwise *Clarkia* interactions were significantly different from zero (no effect of increasing density). Finally, we estimated pollen limitation to successful seed set of each *Clarkia* species as the difference in the probability of successful seed set between pollen addition vs. no addition treatments.

### Total fecundity

The second analysis focused on the total fecundity (total seed number) of focal plants in the interaction plot. We calculated total fecundity as the product of fruit number on a plant, the average number of ovules, seeds, and aborted seeds across the two sampled fruits (control and pollen-supplemented), and the proportion of successful seeds in fruits under each pollination treatment. This allowed us to calculate total estimated reproductive output of the whole plant under the two treatment scenarios. We used generalized linear mixed effects models with a Poisson error distribution to model total reproductive output with six potential fixed effects: pollination treatment, focal *Clarkia* species identity, background *Clarkia* species identity, background *Clarkia* biomass, and the number of forbs and number of grasses within 5cm of the focal seedling. Forb and grass counts were centered and standardized to improve model performance. We included two random effects: an observation-level random effect to account for over-dispersion in the data, and plant identity. We again did not include random effects for plot and block because such a model was unidentifiable.

As before, we built a set of candidate models relevant to the study with which to perform model selection. The most complex candidate model included an interaction between the pollination treatment, background *Clarkia* identity, focal *Clarkia* identity, and background *Clarkia* biomass, as well as the additive effects of forb and grasses (Table S3). This candidate model represented our prediction that whole-plant seed production depends on the density dependent effect of the background *Clarkia* on the focal *Clarkia*, which is modified by the activity of pollinators. We again selected the model with the lowest AICc score for ecological inference. With the best-fit model, we used the Wald test to determine the significance of the predictors on total reproductive output.

### Pollinator preference in the pairwise interaction plots

We built an interaction network using the R package bipartite to understand pollinator preference (Dormann et al. 2008). In the network, we only included the first visit of each pollinator to a plot because each plot was dominated by a single background species such that subsequent movements within a plot were likely to be to the same species (visits to focal plants in plots against a common background species were rare). We first estimated d’, or network specialization, of the three most common pollinator taxa (Blüthgen et al. 2006). Higher values of d’ indicate higher values of pollinator specialization within the network, which we interpret as taxon-level pollinator preferences for different *Clarkia*. We only included the three most common pollinator taxa because all other pollinators accounted less than 2% of the sample (Table S2).

Next, we used a chi-squared test to compare the proportion of first-choice visits to each *Clarkia* species with the proportion of each background plot type we observed. If bees have no preference among *Clarkia*, then the proportion of first-choice visits to each plant species should be equal to the proportion of plots we observed where that species was sown into the background.

### Pollinator constancy in the experimental arrays

To understand the potential for pollinator (in)constancy to affect pairwise interactions between species, we built a 4x4 matrix for each array run, where rows represent the *Clarkia* species first visited by a pollinator, and columns represent the species visited next in a sequence of two sequential flower visits. On-diagonal entries in the matrix represent pollinator movements to the same species of *Clarkia* (constancy), and off-diagonal movements between different species (inconstancy). We divided the sum of all inconstant movements by the total number of moves in arrays to quantify the proportion inconstant movements that might lead to incompatible pollen transfer in *Clarkia*. We also tabulated the number of inconstant movements between each pair of *Clarkia* species and conducted a chi-squared test to determine which pairs of species are at higher risk for incompatible pollen transfer.

### Joint attraction in the pairwise interaction plots

We used a generalized linear mixed effects model of visitation rate (visits per 10-minute observation interval) with a Poisson error distribution to determine the potential for joint attraction in the pairwise interaction plots. Log-transformed floral abundance was used as a fixed effect in the model, and we included random effects for plot and observation period. Joint attraction in this analysis would be an increase in visitation rate as a function of floral abundance.

## Results

### The effect of interactions during growth and flowering on seed production

We collected 371 fruits from 232 focal *Clarkia* plants in the pairwise interaction plots. The uneven number of fruits collected was due to fruit loss from herbivory. We found that the best-fit model of the probability of successful seed set included the four-way interaction between focal *Clarkia* species identity, background *Clarkia* species identity, background *Clarkia* biomass, and pollination treatment (Table S3). In other words, the best fit model of the probability of seed set depended on the species-specific, density dependent effect of background *Clarkia* on focal *Clarkia*, and this effect was modified by pollination treatments.

Pollination significantly changed the density dependent effect of background *Clarkia* on focal *Clarkia* seed set in 31% of the interactions (five of 16 interactions: Figure 2, Table 1). In three of these interactions, the density dependent effect was more competitive when the effect of pollinators was included (no pollen addition treatment) than when the effect of pollinators was removed (pollen addition treatment). In other words, pollinator-mediated interactions introduced a competitive effect of the background *Clarkia* on focal *Clarkia* for the following three background/focal species combinations: *C. unguiculata/C. xantiana; C. xantiana/C. unguiculata;* and *C. cylindrica/C. xantiana*. Conversely, in the other two cases, the density dependent effect of the background was more positive with the effect of pollinators than without; that is, pollinator-mediated interactions introduced a facilitative effect of the background on focal *Clarkia* in *C. speciosa/C. cylindrica* and *C. speciosa/C. speciosa* background/focal combinations. Of these interactions, only two of the 32 density dependent effects of background *Clarkia* density on *Clarkia* seed set were significantly different from zero. One was the competitive effect of *C. unguiculata* on *C. xantiana* in the pollen addition treatment (z =-2.49, p=0.01), and the other was the competitive effect of *C. cylindrica* on *C. xantiana* in the no pollen addition treatment (z= - 4.89, p<0.001) (Figure 2).

**Figure 2.**
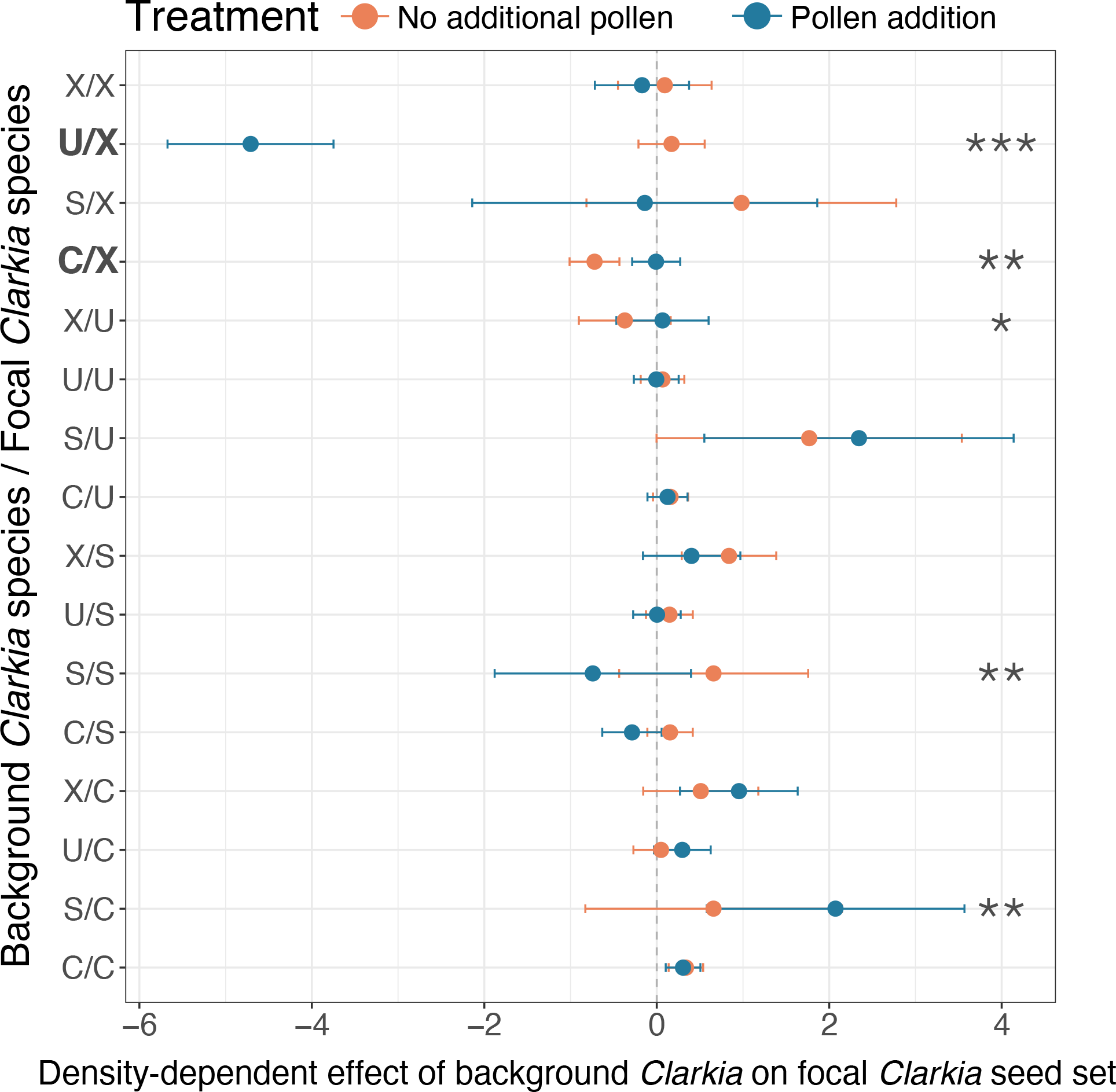
The effect of increasing density of background *Clarkia* on the probability of an ovule becoming a successful seed in focal *Clarkia*. The effect (slope) varies according to species pairs and interactions over pollination. Each row is a different background/focal species combination. Points indicate the estimated effect of increasing background species’ density on the probability of seed set for the focal species, and lines are standard errors. Negative values indicate the probability of seed set declines with increasing background density (a competitive effect of background *Clarkia*), whereas positive values mean that the probability of seed set increases with background density (there is a facilitative effect of background *Clarkia*). Letters stand for the first letter of the species name: C- *Clarkia cylindrica*; S- *C. speciosa*; U-*C. unguiculata*; and X- *C. xantiana*. Blue points are the treatment where the effect of pollinators is removed (pollen addition). Orange points are the treatment where the effect of pollinators is retained (no additional pollen). Bolded combinations are those that include an interaction between background density and pollination treatment that is significantly different from zero; stars indicate a significant effect of pollination treatment on the interaction (*, p<0.05; **, p<0.01; ***, p<0.001).

**Table 1.** The effect of pollinators on seed set in the density dependent interactions between *Clarkia* species (Figure 1), and the suggested explanation of the change in terms of observed pollinator behaviors (Figure 3). Only statistically significant pairwise interactions are shown in this table.

| Background | Focal | Change in density<br>dependent effect<br>of background on<br>focal ± SE | Interpretation | Behavioral<br>Explanation | p-value |
| --- | --- | --- | --- | --- | --- |
| *Clarkia*<br>*unguiculata*<br>(U) | *Clarkia*<br>*xantiana*<br>(X) | 4.88 ± 0.93 | Pollinators<br>alleviate the<br>competitive<br>effect of U on X | Joint attraction to<br>higher densities,<br>but pollinator<br>preference for *C.*<br>*xantiana* | <<br>0.001<br>(***) |
| *Clarkia*<br>*cylindrica*<br>(C) | *Clarkia*<br>*xantiana*<br>(X) | -0.72 ± 0.21 | Pollinators<br>exacerbate the<br>competitive<br>effect of C on X | Incompatible<br>pollen transfer | 0.0011<br>(**) |
| *Clarkia*<br>*xantiana*<br>(X) | *Clarkia*<br>*unguiculata*<br>(U) | -0.44 ± 0.21 | Pollinators<br>exacerbate the<br>competitive<br>effect of X on U | Pollinator<br>preference for *C.*<br>*xantiana* and<br>incompatible<br>pollen transfer | 0.039<br>(*) |
| *Clarkia speciosa*<br>(S) | *Clarkia speciosa*<br>(S) | 1.40 ± 1.48 | Pollinators<br>alleviate the<br>competitive<br>effect of S on S | Pollinator<br>attraction to higher<br>densities | 0.0091<br>(**) |
| *Clarkia speciosa*<br>(S) | *Clarkia cylindrica*<br>(C) | -1.41 ± 0.53 | Pollinators<br>reduce the<br>facilitative effect<br>of S on C | Low pollinator<br>overlap and<br>therefore low<br>pollinator service<br>to C | 0.0073<br>(**) |

Finally, the average pollen limitation to successful seed set varied by focal species. Additional pollen increased seed set in three of the four species: *C. unguiculata, C. cylindrica,* and *C. xantiana*. The treatment effect was highest for *C. unguiculata*, indicating that pollination service limited its successful seed set the most of the four species (Figure 3, Table S4). The treatment effect was lowest in *C. speciosa*, which experienced slightly lower seed success under pollen supplementation (Figure 3, Table S4).

**Figure 3.**
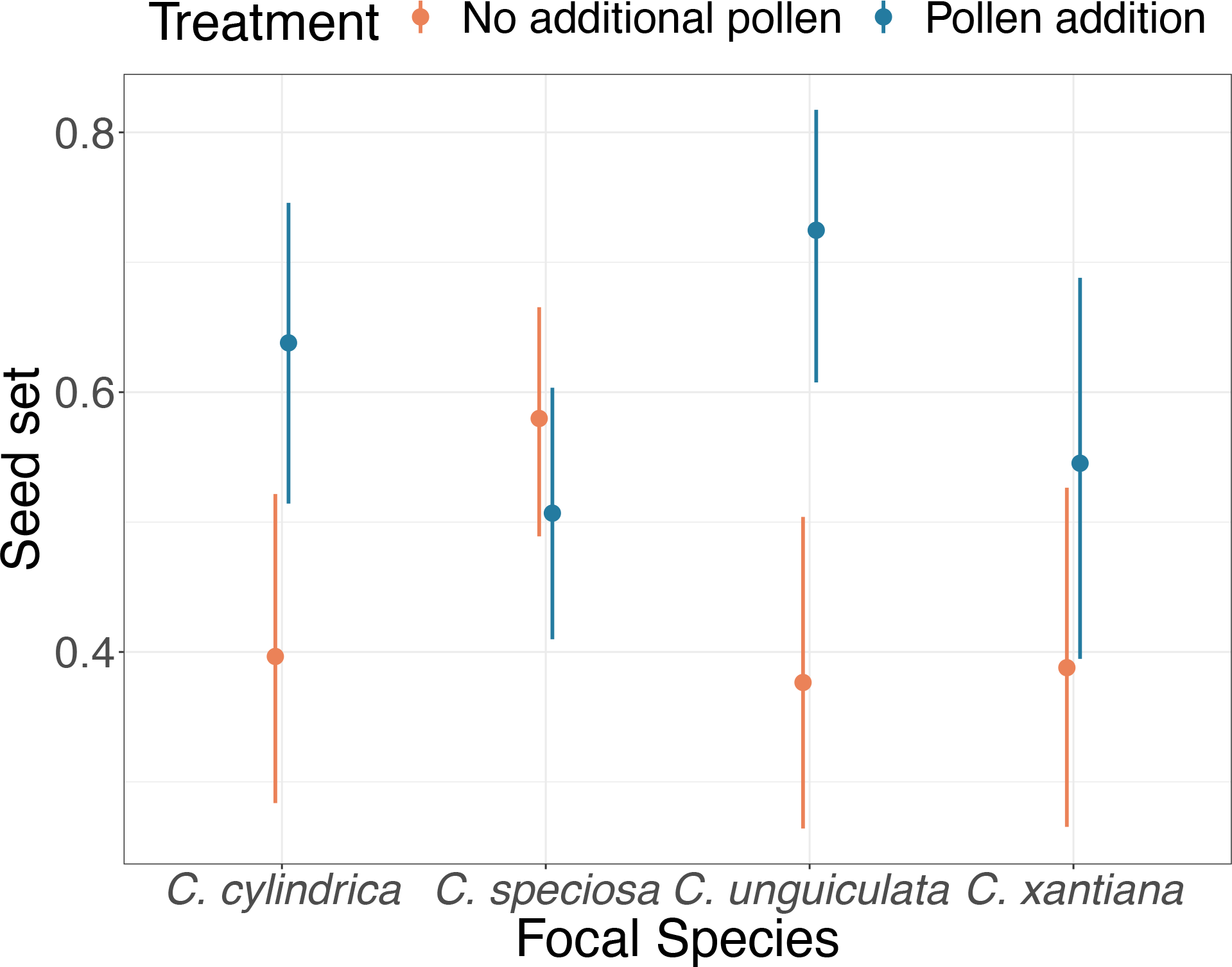
Species-specific probability of seed success in fruits for both pollination treatments. Points represent the marginal estimated average across all background species treatments at the mean background *Clarkia* density. Bars represent the 95% confidence interval for each estimate.

The best-fit model for total fecundity included the additive fixed effects of (1) pollination treatment and (2) the number of forbs surrounding the focal plant when it was a seedling (Table S2). In other words, total fecundity of focal *Clarkia* plants was best predicted by the effect that non-*Clarkia* forbs had on focal plants during growth, and wholesale pollen limitation in *Clarkia* during flowering (Figure 4). Total fecundity strongly declined with increasing forbs (-0.77 ± 0.15 SE; Wald p-value < 0.001, Table S5) and was significantly higher in the pollen addition treatment relative to the no pollen addition treatment (1.17 ± 0.20 SE; Wald p-value < 0.001, Table S5).

**Figure 4.**
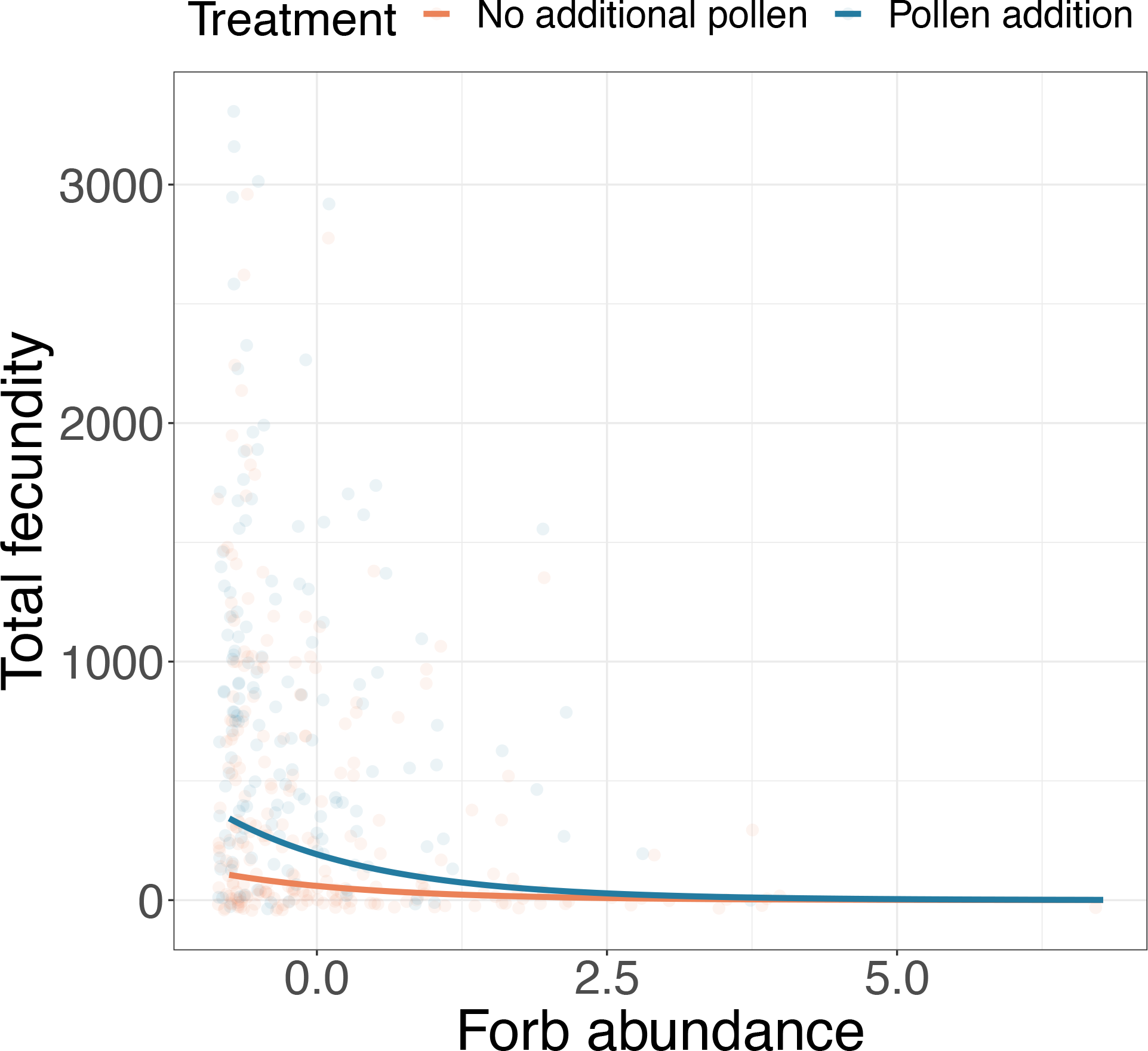
Predicted full-plant seed set as a function of pollination treatment and the number of non-*Clarkia* forbs surrounding the focal as a seedling.

### Pollinator foraging behaviors

#### Preference and joint attraction in the interaction plots

We performed 193, 10-minute observation periods and observed 446 pollinators in the *Clarkia* interaction plots in 2017. The three most abundant pollinator visitors, in order of abundance, were *Diadasia angusticeps* (Apidae), *Hesperapis regularis* (Mellitidae), and *Lasioglossum spp.* (Halictidae) bees, which made up more than 99% of all observed pollinators (Table S2). The preference of these three pollinators varied: *D. anguisticeps* exhibited the most specialized behavior, visiting *Clarkia speciosa* to the near exclusion of the other *Clarkia* (d’ = 0.79), followed by *H. regularis* (d’=0.33), and *Lasioglossum sp.* (d’=17), both of which visited all four *Clarkia*, but most commonly *C. xantiana* and *C. cylindrica* (Figure 5, Panel A).

**Figure 5.**
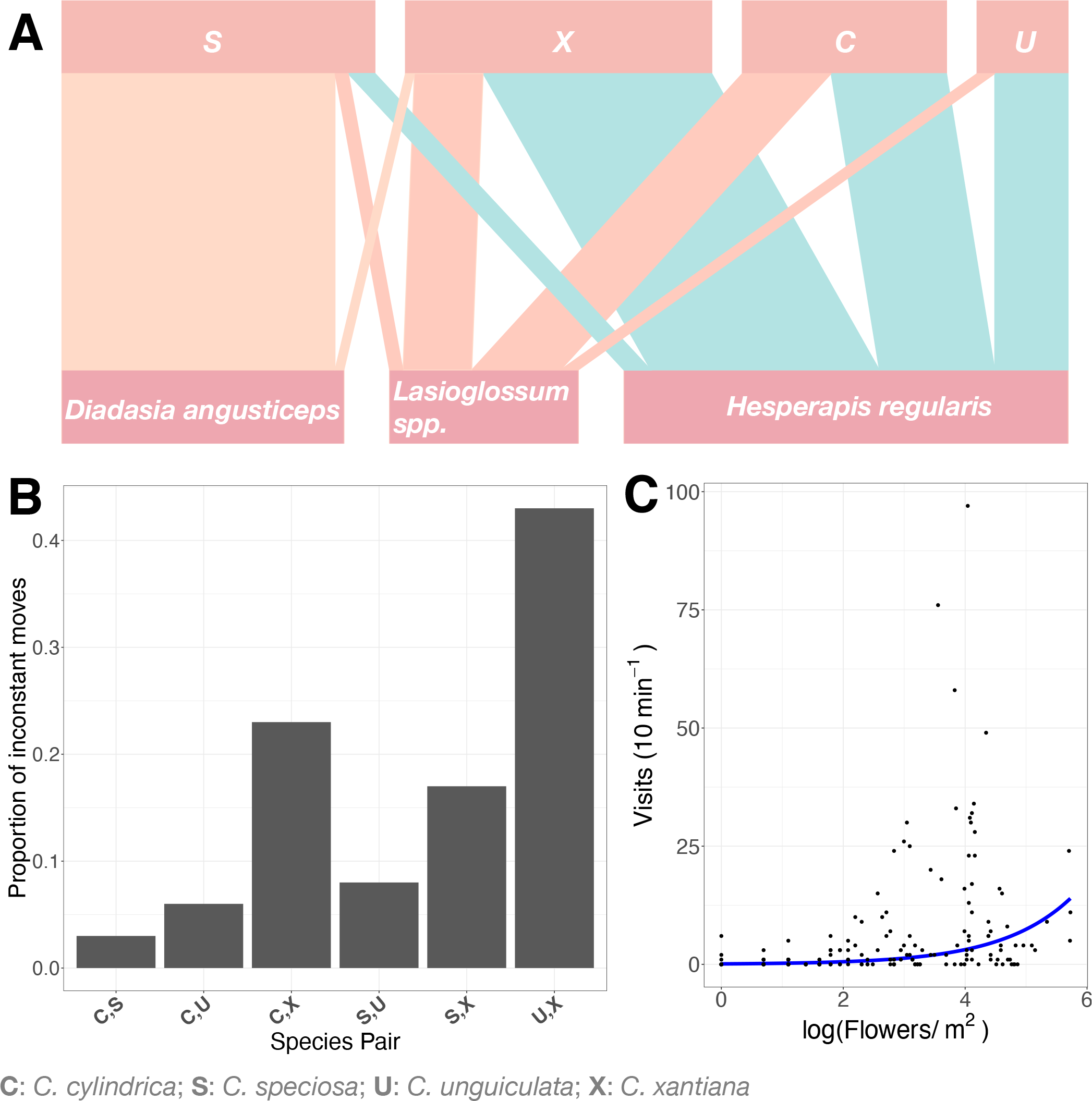
(A) The network of the plant and pollinator interactions observed during the pairwise interaction experiment. (B) The proportion of inconstant moves for each transition type between different *Clarkia* in the experimental arrays. (C) Rate of visitation as a function of floral abundance in the interaction plots.

Overall visitation to the different *Clarkia* was significantly different from the null expectation that visitation to each species is proportional to the frequency with which we observed plots with each background-sown *Clarkia* (χ^2^(3)=104.47; p<0.001). The residuals of the chi-squared analysis indicate pollinator preference for *C. speciosa* and *C. xantiana* (Pearson residuals = 7.6 and 2.4, respectively), and avoidance of *C. cylindrica* and *C. unguiculata* (Pearson residuals = - 3.7, -5.2, respectively).

### Pollinator constancy in the experimental arrays

We performed 17 array runs with all four *Clarkia* species, observing 664 pollinator movements from 418 bees across 2015 and 2016. Only 100 of the 664 movements (15%) were between different plant species (and thus inconstant); therefore, *Clarkia* pollinators are by and large constant foragers, even when equally exposed to all four *Clarkia* species. Interspecific movements were significantly different from the null expectation of being evenly divided among species pairs (χ^2^(5)=65.56; p<0.001). The species pairs where switching occurred most frequently were *C. xantiana* and *C. unguiculata* and *C. xantiana* and *C. cylindrica*. These were the only species pairs with Pearson residuals > 1 (Pearson residuals=6.45 and 1.55 respectively). Movements between individuals in these pairs of accounted for more than 60% of inconstant movements in the dataset (Figure 5, Panel B).

### Pollinator Joint Attraction

Pollinator visitation rate (in visits per 10-minute observation periods) increased with log floral abundance in the interaction plots, indicating joint attraction (0.86±0.11, p<0.001; Figure 5, Panel C).

## Discussion

In community ecology, density dependent species interactions are the building blocks of theories of community function and species persistence. In focusing on either the combined effects of interaction types on species interactions or a single interaction type, studies often overlook the multiplex nature of species interactions which can result in overly simplistic predictions about plant community dynamics (Losapio et al. 2019, Bartomeus et al. 2021). In this study we do our best to reconcile that plant interactions during both growth and flowering can affect density dependent interactions between co-flowering plant species in a group of four sympatric pollinator-sharing *Clarkia* species.

### The importance of studying multiple life history stages to understand plant interactions

We found that the contribution of pollinator-mediated interactions during flowering to total plant fecundity is, above all, contingent on what transpires during growth. Whereas successful seed set depended on pollinator-mediated *Clarkia* interactions (discussed below), the only density dependent interaction explaining the total fecundity of focal plants was that of the non-*Clarkia* forbs surrounding seedlings during growth. We anticipated that the effect of interactions during flowering might change the direction and/or magnitude of density dependence independently of the effects of growing together, which is how previous studies have typically framed the question of how pollinators could affect plant coexistence dynamics (e.g. Benadi & Pauw 2018, Bergamo et al. 2020, Lanuza et al. 2018, Lázaro et al. 2014, Rathcke 1983). In contrast, our findings were surprising in two ways: first, the competitive effect exerted by plants during growth limited total fecundity so strongly, by limiting total ovule production, that it masked the effect of pollinator-mediated interactions on seed production per ovule during flowering. This demonstrates that the significance of pollinator-mediated plant interactions for plant fecundity is constrained by, rather than independent of, plant interactions during growth.

Second, density dependent competition during growth was exerted by non-*Clarkia* forbs on *Clarkia* seedlings, whereas the density dependent effects of pollinator-mediated interactions during flowering were exerted by *Clarkia* on *Clarkia*. This result demonstrates that the density dependent effects of particular species (in this case, congeneric *Clarkia*) are potentially only relevant during one, rather than all, life stages (Leslie 2005). Stage-dependent interactions have been explored in animal communities in terms of ontogenetic niche shifts (Werner & Gillam 1984, Bassar et al. 2017), and our results suggest that accounting for ontogeny may be appropriate for understanding plant community coexistence as well. This result also highlighted that studying *Clarkia* interactions without removing background grasses and forbs was revealing of the ecology of the plant community. Had we taken a more traditional approach of studying pairwise interactions without the rest of the annual plant community (by weeding, for example), we may have concluded that interactions during flowering are strong drivers of *Clarkia* interaction dynamics, because it would have been impossible to detect the density dependent effects introduced by non-*Clarkia* forbs. Our results support the idea that studying species interactions in a pairwise fashion can limit our understanding of community structure and function (Abrams 1983, Chesson 2018, Levine et al. 2017, Mayfield & Stouffer 2017). One way to incorporate such complexity into predictive models may be to leverage the growing body of network theory (Godoy et al. 2018, Opedal & Heglend 2020, Valdovinos 2019, Losapio et al. 2019) to build models that capture how the stage-specificity and contingency of density dependent species interactions affect population persistence and community structure. Alternatively, the implications of our findings could be explored by using population models to compare plant species persistence in communities where interactions during growth take precedence in determining fecundity, versus communities where pollinator-mediated interactions determine fecundity.

### Pollinator-mediated interactions correspond to foraging behavior and affect seed set

We have shown that pollinator-mediated interactions during flowering affect seed set at the scale of the fruit and can be linked to the foraging behaviors of pollinators. This links two types of pollination ecology: studies that show that pollinator foraging varies according to community context (Eckhart et al. 2006, Lázaro et al. 2009, Valdovinos et al. 2013), and that pollinator sharing among species affects focal plant performance (Bergamo et al. 2019, Moeller 2004, Lanuza et al. 2018). Here we found that pollinators changed the positive, negative, or neutral effect of background *Clarkia* density on seed set in five of 16 species interactions. A few patterns stand out from this analysis. First, the *Clarkia* pairs that exhibited density dependent interactions significantly different than zero (*C. cylindrica* background*/C. xantiana* focal, and *C. unguiculata* background*/C. xantiana* focal) shared pollinators and represented the two species pairs where pollinators were most likely to make interspecific (inconstant) visits (Figure 5, Panel B). In one of these interactions, the significantly competitive effect of *C. unguiculata* density on *C. xantiana* was neutralized when including the effect of pollinator sharing. To wit, pollinators acted as a facilitative force and ‘rescued’ *C. xantiana* from the otherwise competitive effect of increasing *C. unguiculata* density. This result may be explained by joint attraction, where increasing *Clarkia* density in the plot attracted more pollinators overall, and that pollinators preferred *C. xantiana* much more than they did *C. unguiculata*, resulting in *C. xantiana* receiving a disproportionate amount of pollinator visits. In the other species interaction significantly different from zero, pollinators introduced a competitive effect of *C. cylindrica* on *C. xantiana*. We suspect this can be explained simply by incompatible pollen transfer between plants from inconstant shared pollinators. Alternatively, the fact that pollinators alleviated competition in one case but exacerbated competition in another could be explained by potential differences in how different species tolerate incompatible pollen. For example, previous work has shown that flowering plants, including *Clarkia*, exhibit species-specific sensitivity or tolerance to heterospecific pollens (Arceo-Gómez et al. 2016, Arceo-Gómez et al. 2019); it is possible that *C. xantiana* tolerates *C. unguiculata* pollen better than it does *C. cylindrica* pollen, resulting in the differences in pollinator-mediated interactions.

In addition, the only species that exhibited a significant intraspecific pollinator-mediated interaction was *C. speciosa,* which does not share pollinators to a great extent with the other three species and experienced lower rates of inconstant visitation. In this instance, increasing conspecific density trended toward competition when the effect of pollinators was removed but trended toward facilitation when the effect of pollinators was retained. This suggests that joint attraction can introduce a positive density dependent Allee effect in conspecific stands of flowering plants, as suggested by a previous model exploring the relationship between pollinator foraging, plant density, and plant performance (Kunin & Iwasa 1996). However, we would likely not have found positive density dependence if not for *Diadasia angusticeps*, which has a decided preference for *C*. *speciosa*, and was quite abundant in 2017. Without *D. anguisticeps*, or in years of low abundance, pollination service to *C. speciosa* would likely have been lower in both quantity and quality both because generalists can exacerbate negative reproductive outcomes (Kunin 1993) and because competition for pollinator services generally increases in plant communities when pollinators are scarcer (Lázaro et al. 2014, Ye et al. 2014).

Finally, the effect of *C. speciosa* backgrounds on *C. cylindrica* seed set trended towards facilitative without the effect of pollinators but became less so with effect of pollinators; in this case, pollinators introduced a competitive effect that dampened *C. speciosa* facilitation of *C. cylindrica*. Because these two species do not share pollinators (Figure 4 Panel A), and there is rarely switching between the two (Figure 4 Panel C), the competitive effect from pollinators is likely due to low pollinator service to *C. cylindrica* when surrounded by *C. speciosa* (Table 1).

### Plant-pollinator networks are predictive of plant average pollen limitation

In our study, network information corresponded to patterns in plant species’ average pollen limitation. Whereas additional compatible pollen did not increase seed set in focal *C. speciosa*, it did increase the average seed set in the three other species: *C. xantiana* experienced a small boost in seed set with additional pollen, *C. cylindrica* a slightly higher boost, and *C. unguiculata* saw the greatest effect of additional pollen on seed set. Thus, high pollinator preference and low rates of pollinator sharing translated to high rates of compatible pollen transfer for *C. speciosa*. In contrast, higher rates of pollinator sharing likely resulted in interspecific pollen transfer in the three other *Clarkia* species resulting in pollen limitation, and the magnitude of pollen limitation corresponded to pollinator preference. The promising implication of this result is that comparatively easy-to-collect information from plant-pollinator interaction networks maps closely to average pollen limitation in plant populations, which is more difficult to measure. Testing this idea in high diversity plant-pollinator systems would help illuminate the potential relationship between plant population level pollen limitation and plant-pollinator networks.

## Conclusion

In this work, we have endeavored to understand how density dependence during growth and flowering, separately and in combination, determine plant seed production. We found that the effects of species interactions in the flowering phase of life were explained by pollinator foraging behaviors, but that these effects on total fecundity were overridden by interactions in the growth phase. In total, this study shows how natural history work describing the complexity of biological interactions can be linked to performance variables relevant to population and community dynamics in ways that are revealing of community function. It may also help frame future theoretical studies of stage-dependent multispecies interactions, as the larger task of identifying the population and community-level ramifications of such interactions remains.

## Literature Cited

Abrams, P.A. 1983. Arguments in Favor of Higher Order Interactions. The American Naturalist 121: 887–891.

Angert, A., Huxman, T., Chesson, P., Venable, D. 2009. Functional Tradeoffs Determine Species Coexistence via the Storage Effect. Proceedings of the National Academy of Sciences 106: 11641–11645.

Arceo-Gómez, G., Raguso, R.A., and Geber, M.A. 2016. Can Plants Evolve Tolerance Mechanisms to Heterospecific Pollen Effects? An Experimental Test of the Adaptive Potential in *Clarkia* Species. Oikos 125: 718–25.

Arceo-Gómez, G., Schroeder, A., Albor, C., Ashman, T.L., Knight, T.M., Bennett, J.M., Suarez, B., and Parra-Tabla, V. 2019. Global Geographic Patterns of Heterospecific Pollen Receipt Help Uncover Potential Ecological and Evolutionary Impacts across Plant Communities Worldwide. Scientific Reports 9:8086.

Barrios, B., Pena, S.R., Salas, A., and Koptur, S. 2016. Butterflies Visit More Frequently, but Bees Are Better Pollinators: The Importance of Mouthpart Dimensions in Effective Pollen Removal and Deposition. AoB PLANTS 8:plw001.

Bartomeus, I., Saavedra, S., Rohr, R.P., Godoy, O. 2021. Experimental Evidence of the Importance of Multitrophic Structure for Species Persistence. Proceedings of the National Academy of Sciences, 118: e2023872118.

Bassar, D.B., Travis, J., and Coulson, T. 2017. Predicting Coexistence in Species with Continuous Ontogenetic Niche Shifts and Competitive Asymmetry. Ecology 98: 2823–2836.

Bates, D., Maechler, M., Bolker, B., Walker, S. 2015. Fitting Linear Mixed-Effects Models Using lme4. Journal of Statistical Software, 671: 1–48.

Benadi, G. 2015. Requirements For Plant Coexistence Through Pollination Niche Partitioning. Proceedings of the Royal Society B 282:20150117.

Benadi, G., and Pauw, A. 2018. Frequency Dependence of Pollinator Visitation Rates Suggests That Pollination Niches Can Allow Plant Species Coexistence. Journal of Ecology 106: 1892– 1901.

Bergamo, P.J., Streher, N.S., Traveset, A., Wolowski, M., and Sazima, M. 2020. Pollination Outcomes Reveal Negative Density Dependence Coupled with Interspecific Facilitation among Plants. Ecology Letters 23:129–139.

Bimler, M. D., Stouffer, D., Lai, H. R., and Mayfield, M.M. 2018. Accurate Predictions of Coexistence in Natural Systems Require the Inclusion of Facilitative Interactions and Environmental Dependency. Journal of Ecology 106: 1839–1852.

Bizecki Robson, D. 2013. An Assessment of the Potential for Pollination Facilitation of a Rare Plant by Common Plants: *Symphyotrichum sericeum* (Asteraceae) as a Case Study. Botany 91: 34–42.

Blüthgen, N., Menzel, F., and Blüthgen, N. 2006. Measuring Specialization in Species Interaction Networks. BMC Ecology 6:9.

Chesson, P. 2018. Updates on Mechanisms of Maintenance of Species Diversity. Journal of Ecology 106: 1773–1794.

Dormann, C.F., Fründ, J., Blüthgen, N. & Gruber B. 2009. Indices, graphs and null models: analyzing bipartite ecological networks. The Open Ecology Journal 2: 7–24.

Dybzinski R., and Tilman, D. 2007. Resource Use Patterns Predict Long-Term Outcomes of Plant Competition for Nutrients and Light. The American Naturalist 170; 305–318.

Eckhart, V., Rushing, N.S., Hart G.M., and Hansen, J.D. 2006. Frequency-Dependent Pollinator Foraging in Polymorphic *Clarkia xantiana* ssp. *xantiana* Populations: Implications for Flower Colour Evolution and Pollinator Interactions. Oikos 112: 412–21.

Eisen, K.E., and Geber, M.A. 2018. Ecological Sorting and Character Displacement Contribute to the Structure of Communities of *Clarkia* Species. Journal of Evolutionary Biology 31: 1440– 58.

Eisen, K.E., Wruck, A.C. and Geber, M. A. 2020. Floral Density and Co-occurring Congeners Alter Patterns of Selection in Annual Plant Communities. Evolution 74:1682–1698.

Feldman, T.S., Morris, W.F. and Wilson, W.G. 2004. When Can Two Plant Species Facilitate Each Other ’s Pollination? Oikos 105:197–207.

Godoy, O., and Levine, J.M. 2014. Phenology Effects on Invasion Success: Insights from Coupling Field Experiments to Coexistence Theory. Ecology 95: 726–36.

Godoy, O., Kraft, N. J. B., and Levine, J. M. 2014. Phylogenetic Relatedness and the Determinants of Competitive Outcomes. Ecology Letters 17:836–844.

Godoy, O., Bartomeus, I., Rohr, R.P., and Saavedra, S. 2018. Towards the Integration of Niche and Network Theories. Trends in Ecology and Evolution 33:287–300.

Gould, B., Moeller, D.A., Eckhart, V.M., Tiffin, P., Fabio, E., and Geber, M. A. 2014. Local Adaptation and Range Boundary Formation in Response to Complex Environmental Gradients Across the Geographic Range of *Clarkia xantiana* ssp. *xantiana*. Journal of Ecology 102:95–107.

Hart, G.M., and Eckhart, V.M. 2010. Ecological Separation in Foraging Schedule and Food Type Between Pollinators of The California Wildflower, Clarkia xantiana ssp. xantiana. Journal of Pollination Ecology 2:13–20.

Hove, A. A., Mazer, S.J., Ivey, C.T. 2016. Seed Set Variation in Wild *Clarkia* Populations: Teasing Apart the Effects of Seasonal Resource Depletion, Pollen Quality, and Pollen Quality. Ecology and Evolution 6: 6524–6536.

Kraft, N., Godoy, O., and Levine, J. 2014. Plant Functional Traits and the Multidimensional Nature of Species Coexistence. Proceedings of the National Academy of Sciences 112: 797–802.

Kunin, W. 1993. Sex and the Single Mustard: Population Density and Pollinator Behavior Effects on Seed Set. Ecology 74: 2145–2160.

Kunin, W., and Iwasa, Y. 1996. Pollinator Foraging Strategies in Mixed Floral Arrays: Density Effects and Floral Constancy. Theoretical Population Biology 49: 232–63.

Lanuza, J. B., Bartomeus, I. and Godoy, O. 2018. Opposing Effects of Floral Visitors and Soil Conditions on the Determinants of Competitive Outcomes Maintain Species Diversity in Heterogeneous Landscapes. Ecology Letters 21: 865–74.

Lázaro, A., Lundgren, R., Totland, Ø., Lázaro, A., Lundgren, R., & Totland, Ø. (n.d.). Co- flowering neighbors influence the diversity and identity of pollinator groups visiting plant species. Oikos 118: 691–702.

Lázaro, A., Lundgren, R., and Totland, Ø. 2014. Experimental Reduction of Pollinator Visitation Modifies Plant–Plant Interactions for Pollination. Oikos 123: 1037–48.

Lenth, R. 2020. emmeans: Estimated Marginal Means, aka Least-Squares Means. R package version 1.4.4. https://CRAN.R-project.org/package=emmeans

Leslie, H. 2005. Positive Intraspecific Effects Trump Negative Effects in High-Density Barnacle Aggregations. Ecology 86: 2716–2725.

Levine, J.M., Bascompte, J., Adler, P.B., and Allesina, S. 2017. Beyond Pairwise Mechanisms of Species Coexistence in Complex Communities. Nature 546: 56–64.

Lewis, H. and Lewis, M. E. 1955. The Genus Clarkia. University of California Publications in Botany 20: 241–392.

Losapio, G., Fortuna, M.A., Bascompte, J. Schmid, B., Michalet, R., Neumeyer, R., Castro, L., Cerretti, P., Germann, C., Haenni, J.P., Klopfstein, S., Ortize-Sanchez, F.J., Pont, A.C., Rousse, P., Schmid, K. Sommaggio, D., and Schöb, C. 2019. Plant Interactions Shape Pollination Networks via Nonadditive Effect. Ecology 100:e02619.

MacSwain, J. W., Raven, P. H. and Thorp, R. W. 1973. Comparative Behaviour of Bees and Onagraceae. IV. Clarkia bees of the western United States. University of Calif. Publ. Entomol. 1:80.

Mayfield, M.M., and Stouffer, D.B. 2017. Higher-order Interactions Capture Unexplained Complexity in Diverse Communities. Nature Ecology and Evolution 1:62.

Moeller, D.A. 2004. Facilitative Interactions among Plants via Shared Pollinators. Ecology 85: 3289–3301.

Moeller, D.A. 2005. Pollinator Community Structure and Sources of Spatial Variation In Plant- Pollinator Interactions In Clarkia xantiana ssp. xantiana. Oecologia 142: 28–37.

Moeller, D.A., Geber, M.A., Eckhart V., and Tiffin, P. 2012. Reduced Pollinator Service and Elevated Pollen Limitation at the Geographic Range Limit of an Annual Plant. Ecology 93: 1036–1048.

Morales, C.L., and Traveset, A. 2008. Interspecific Pollen Transfer: Magnitude, Prevalence and Consequences for Plant Fitness. Critical Reviews in Plant Sciences, 27:221–238.

Moreira-Hernández, J.I., and Muchhala, N. 2019. Importance of Pollinator Mediated Interspecific Pollen Transfer for Angiosperm Evolution. Annual Review of Ecology, Evolution, and Systematics 50:191–217.

Opedal, Ø.H., and Hegland, S.J. 2020. Using Hierarchical Joint Models to Study Reproductive Interactions in Plant Communities. Journal of Ecology 108:485–495.

Pauw, A. 2012. Can Pollination Niches Facilitate Plant Coexistence? Trends in Ecology & Evolution 28:30–37.

Phillips R.D., Peakall R., van der Niet T., and Johnson S.D. 2020. Niche Perspectives on Plant- Pollinator Interactions. Trends in Plant Science 25:779–793.

Rathcke, B. 1983. Competition and Facilitation Among Plants for Pollination. In Leslie Real Editor, Pollination Biology, 305–329.

Singh, I. 2014. Pollination interaction networks between Clarkia (Onagraceae) species and their pollinators in the Southern Sierra Nevada, California. Cornell University.

Song, Z. and Feldman, M.W. 2014. Adaptive Foraging Behaviour of Individual Pollinators and the Coexistence of Co-flowering Plants. Proceedings of the Royal Society B 281: 20132437.

Valdovinos F.S., Moisset de Espanés P., Flores J.D., and Ramos-Jiliberto R. 2013. Adaptive Foraging Allows the Maintenance of Biodiversity of Pollination Networks. Oikos 122: 907–917.

Valdovinos, F. S. 2019. Mutualistic Networks: Moving Closer to a Predictive Theory. Ecology Letters, *22*L 1517–1534.

Wainwright, C., HilleRisLambers, J., Lai, H. Loy, X. and Mayfield. M.M. 2019. Distinct Responses of Niche and Fitness Differences to Water Availability Underlie Variable Coexistence Outcomes in Semi-arid Annual Plant Communities. Journal of Ecology 107: 293–306.

Waser, N.M., Chittka, L., Price, M.V., Williams, N.M., and Ollerton, J. 1996. Generalization in Pollination Systems, and Why It Matters. Ecology 77:1043–60.

Werner, E.E. and Gillam, J.F. 1984. The Ontogenetic Niche and Species Interactions in Size- Structured Populations. Annual Review of Ecology and Systematics 15: 393–425.

Ye, Z.M., Dai, W.K., Jin, X.F., Gituru, R.W., Wang, Q.F., and Yang, C.F. 2014. Competition and Facilitation among Plants for Pollination: Can Pollinator Abundance Shift the Plant-Plant Interactions? Plant Ecology 215:3–13.

