## Supplemental information for "Density dependent interactions during growth mask the effect of interactions during flowering in pollinator-sharing annual plants"

**Supplemental Materials**

Table S1. List of common grasses and forbs in interaction plots at field site.

| Grass (G) or Forb (F) | Genus and species | Family |
| --- | --- | --- |
| G | *Avena barbata* | Poaceae |
| G | *Bromus diandrus* | Poaceae |
| G | *Bromus rubens* | Poaceae |
| F | *Dichelostemma congesta* | Asparagaceae |
| F | *Senecia sp.* | Asteraceae |
| F | *Amsinckia intermedia* | Boraginaceae |
| F | *Cryptantha intermedia* | Boraginaceae |
| F | *Pagiobothrys* spp. | Boraginaceae |
| F | *Phacelia tanacetifolia* | Boraginaceae |
| F | *Phacelia cicutaria* | Boraginaceae |
| F | *Pholistoma auritum* | Boraginaceae |
| F | *Lotus micranthus* | Fabaceae |
| F | *Lupinus bicolor* | Fabaceae |
| F | *Erodium cicutarium* | Geraniaceae |
| F | *Salvia columbariae* | Lamiaceae |
| F | *Claytonia perfoliate* | Montiaceae |
| F | *Castilleja exerta* | Orobanchaceae |
| F | *Gilia* sp. | Polemoniaceae |
| F | *Linanthus dichotomous* | Polemoniaceae |

Table S2. Pollinators observed visiting *Clarkia* plants in the pairwise interaction plot experiment. Bolded rows the taxa we included in the analysis.

| Bee genus | Number of observations | Proportion of total visitors |
| --- | --- | --- |
| ***Hesperapis*** | **200** | **0.45** |
| ***Diadasia*** | **127** | **0.28** |
| ***Lasioglossum*** | **85** | **0.19** |
| *Megachile* | 4 | 0.008 |
| *Agapostemon* | 1 | 0.002 |
| *Apis* | 1 | 0.002 |
| *Eucera* | 1 | 0.002 |
| *Osmia* | 1 | 0.002 |
| *Unknown* | 26 | 0.06 |

| **Variable** | **Family** | **Formula** | **deviance** | **residual df** | **delta AICc** | **AICc Weight** |
| --- | --- | --- | --- | --- | --- | --- |
| **Seed set** | **Binomial** | **probability of seed set ~ pollination treatment * focal species * background species * background *Clarkia* biomass + (1\|plant)** | **3294.7** | **306** | **0** | **3452.871** |
|  |  | probability of seed set ~ pollination treatment * focal species * background species + (1\|plant) | 3436.2 | 338 | 56.019 | 3508.89 |
|  |  | probability of seed set ~ pollination treatment * focal species * background species + background *Clarkia* biomass + (1\|plant) | 3435 | 337 | 57.223 | 3510.094 |
|  |  | probability of seed set ~ pollination treatment * focal species + (1\|plant) | 3590 | 362 | 155.601 | 3608.472 |
|  |  | probability of seed set ~ pollination treatment + focal species + (1\|plant) | 3803.6 | 365 | 362.91 | 3815.781 |
|  |  | probability of seed set ~ pollination treatment + focal species + background *Clarkia* biomass + (1\|plant) | 3803.1 | 364 | 364.553 | 3817.424 |
| **Total fecundity** | **Poisson with OLRE** | **whole plant seed set ~ pollination treatment + background forb count + (1\|plant) + (1\|observation number)** | **5249.9** | **366** | **0** | **5260.054** |
|  |  | whole plant seed set ~ pollination treatment + background grass count + background forb count + (1\|plant) + (1\|observation number) | 5249.9 | 365 | 2.066 | 5262.12 |
|  |  | whole plant seed set ~ pollination treatment + (1\|plant) + (1\|observation number) | 5273.4 | 367 | 21.457 | 5281.511 |
|  |  | whole plant seed set ~ pollination treatment + background Clarkia biomass + (1\|plant) + (1\|observation number) | 5273.4 | 366 | 23.499 | 5283.553 |
|  |  | whole plant seed set ~ pollination treatment * background Clarkia biomass + (1\|plant) + (1\|observation number) | 5272.5 | 365 | 24.686 | 5284.74 |
|  |  | whole plant seed set ~ pollination treatment + focal species + (1\|plant) + (1\|observation number) | 5272.5 | 364 | 26.774 | 5286.828 |
|  |  | whole plant seed set ~ pollination treatment + focal species + background Clarkia biomass + (1\|plant) + (1\|observation number) | 5272.5 | 363 | 28.857 | 5288.911 |
|  |  | whole plant seed set ~ pollination treatment * focal species + (1\|plant) + (1\|observation number) | 5272 | 361 | 32.542 | 5292.596 |
|  |  | whole plant seed set ~ pollination treatment * focal species * background species + (1\|plant) + (1\|observation number) | 5238.7 | 337 | 53.708 | 5313.762 |
|  |  | whole plant seed set ~ 1 + (1\|plant) + (1\|observation number) | 5309 | 368 | 54.984 | 5315.038 |
|  |  | whole plant seed set ~ pollination treatment * focal species * background species + background Clarkia biomass+ (1\|plant) + (1\|observation number) | 5238.6 | 336 | 56.097 | 5316.151 |
|  |  | whole plant seed set~background Clarkia biomass + (1\|plant) + (1\|observation number) | 5309 | 367 | 57.027 | 5317.081 |
|  |  | whole plant seed set ~ focal species + (1\|plant) + (1\|observation number) | 5307.1 | 365 | 59.275 | 5319.329 |
|  |  | whole plant seed set ~ pollination treatment * focal species * background species * background Clarkia biomass + (1\|plant) + (1\|observation number) | 5194.7 | 305 | 95.749 | 5355.803 |

Table S3. Candidate models and model results used in model selection for explaining variation in seed set and whole plant fecundity.

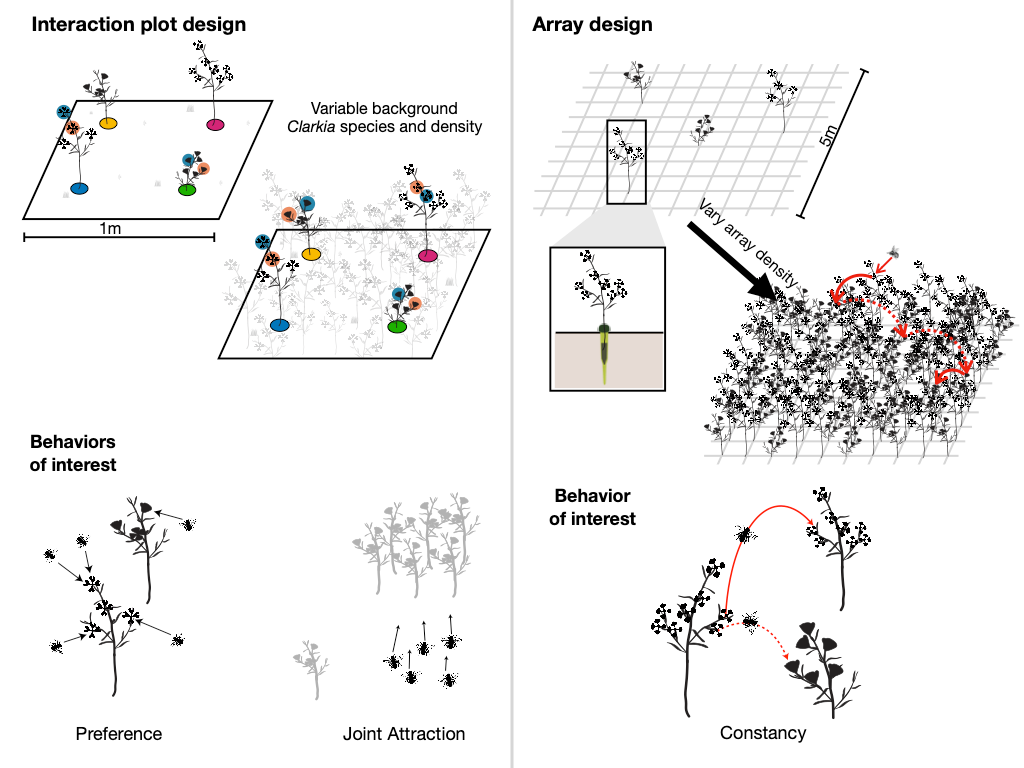

Figure S1. Pollinator behaviors of interest and how we assessed them. In the left-hand panel, we show how we collected information about preference and joint attraction in the interaction plots. In the right-hand panel, we show how we collected information about pollinator constancy using experimental arrays.

Table S4. Change in seed set between treatments and effect size as measured by Cohen’s d. Negative values indicate that seed set increased with additional pollen. *Clarkia speciosa* has a marginally positive value of Cohen’s d, indicating that seed set was higher in control fruits, likely due to high quality pollination service such that additional pollination clogged the stigma of some fruits.

| **Focal Species** | **Treatment** | **Seed set** | **Lower CI (95%)** | **Upper CI (95%)** | **Effect size (Cohen's d) ± SE** |
| --- | --- | --- | --- | --- | --- |
| *C. cylindrica* (C) | No Additional Pollen | 0.40 | 0.28 | 0.52 | -0.986 ± 0.094 |
|  | Additional Pollen | 0.63 | 0.51 | 0.75 |  |
| *C. speciosa* (S) | No Additional Pollen | 0.58 | 0.49 | 0.67 | 0.294 ± 0.106 |
|  | Additional Pollen | 0.51 | 0.41 | 0.60 |  |
| *C. unguiculata* (U) | No Additional Pollen | 0.38 | 0.25 | 0.50 | -1.472 ± 0.121 |
|  | Additional Pollen | 0.72 | 0.61 | 0.82 |  |
| *C. xantiana* (X) | No Additional Pollen | 0.38 | 0.27 | 0.53 | -0.638 ± 0.204 |
|  | Additional Pollen | 0.55 | 0.40 | 0.69 |  |

Table S5. Model coefficients for the best-fit model of total fecundity. Forb abundance in this analysis was centered and standardized.

| *Coefficient* | *Log-Mean* | *Conf. Int (95%)* | *P-Value* |
| --- | --- | --- | --- |
| Control | 4.09 | 3.74 – 4.43 | **<0.001** |
| Supplement | 1.17 | 0.77 – 1.57 | **<0.001** |
| Forb abundance | -0.77 | -1.07 – -0.46 | **<0.001** |
| **Random Effects** |  |  |  |
| σ^2^ | 2.88 |  |  |
| τ_00_ _re_ | 2.87 |  |  |
| τ_00_ _plantid_ | 3.99 |  |  |
| ICC | 0.58 |  |  |
| N _plantid_ | 232 |  |  |
| N _re_ | 371 |  |  |
| Observations | 371 |  |  |
| Marginal R^2^ / Conditional R^2^ | 0.130 / 0.635 | |  |
